## Supporting Information for "Ultrafast phasor-based hyperspectral snapshot microscopy for biomedical imaging"

| | |
|---|---|
| Supplementary Fig. S1 | Schematic of the sideSPIM setup and sine/cosine and bandpass filter spectra measured with an absorption spectrometer. |
| Supplementary Fig. S2 | Fluorescent dye solutions measured with the sine/cosine filter method compared to 32 channel spectral detection. |
| Supplementary Fig. S3 | Raw images and phasor plot histograms of eleven dye solutions and demonstration of the rule of linear addition. |
| Supplementary Fig. S4 | Phasor plot rule of linear addition demonstrated by simulations. |
| Supplementary Fig. S5 | Raw sine, cosine, total intensity, unmixed phasor and bandpass filter images of live cells stained with four dyes. |
| Supplementary Fig. S6 | Single stained NIH 3T3 cells and corresponding phasor plot distributions. |
| Supplementary Fig. S7 | SoFa transgenic 72 hpf zebrafish imaged with conventional bandpass filters and the sine/cosine hyperspectral method. |
| Supplementary Fig. S8 | Hyperspectral cellular fingerprinting reveals zebrafish retinal composition. |
| Supplementary Fig. S9 | Principle of phasor plot analysis based on linear combinations. |
| Supplementary Fig. S10 | Raw sine, cosine, and total intensity images of a SoFa zebrafish retina. |
| Supplementary Fig. S11 | Spectral phasor positions of ACDAN, NADH and FAD in aqueous solutions. |
| Supplementary Movie M1 | 5D rendered time lapse sequence shown in Fig. 3m-q. |
| Supplementary Movie M2 | 360º rotation of the color mapped 3D volume shown in Fig. 3l. |
| Supplementary Movie M3 | 360º rotation of the color mapped 3D volume shown in Fig. 4c. |

**Supplementary Fig. S1.**

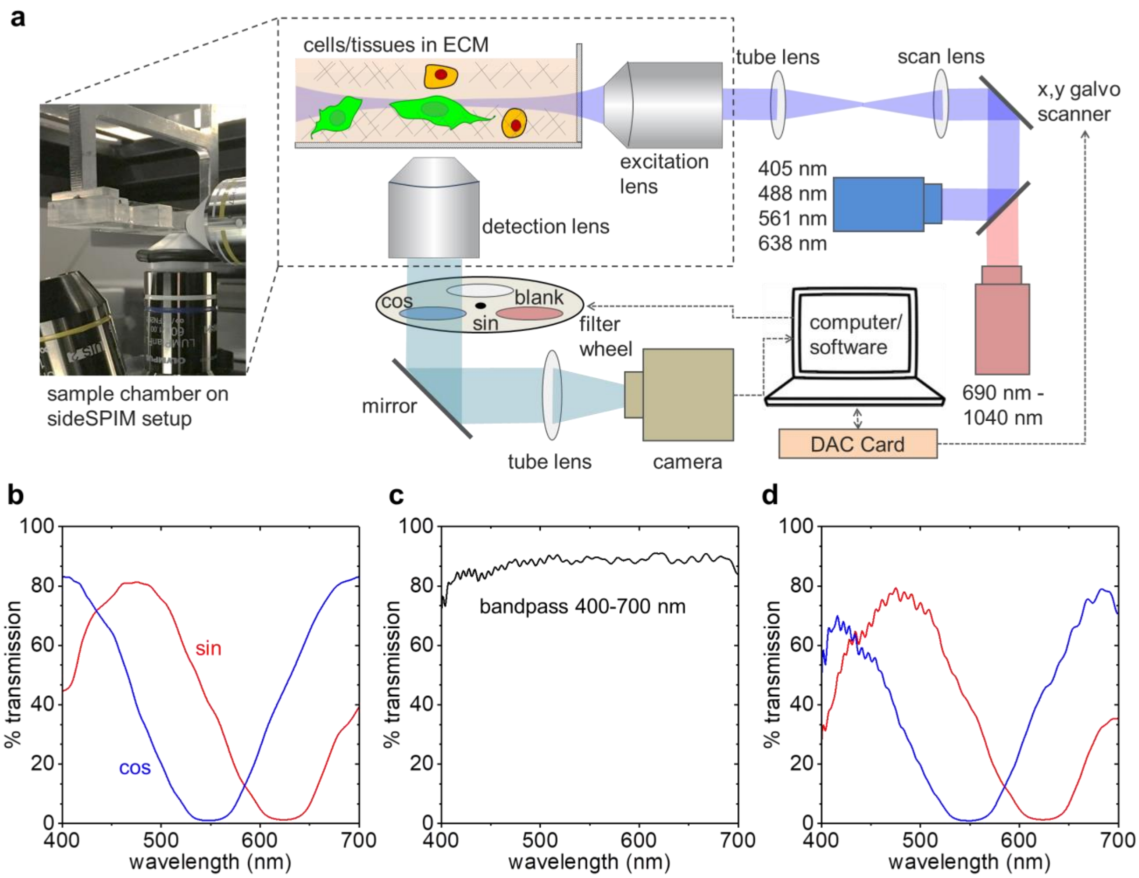


**Schematic of the sideSPIM setup and sine/cosine and bandpass filter spectra measured with an absorption spectrometer.** (**a**) The light sheet microscope was based on our sideSPIM setup as described in the Methods section. (**b**) Absorption spectra of sine/cosine interference filters. (**c**) Absorption spectrum of 400-700 nm bandpass interference filter. (**d**) Combined sine/cosine and 400-700 nm bandpass interference filter absorption spectra. Filters were custom ordered from OptoSigma (Irvine, CA) as specified in the Methods section.

**Supplementary Fig. S2.**

| Dye | G(sin/cos) | G(32ch) | S(sin/cos) | S(32ch) | λ(sin/cos) (nm) | λ(32ch) (nm) | M(sin/cos) | M(32ch) |
|---|---|---|---|---|---|---|---|---|
| Rhodamine 110 | -0.834 | -0.736 | 0.145 | 0.258 | 542 | 538 | 0.847 | 0.780 |
| Rhodamine 6G | -0.779 | -0.726 | -0.394 | -0.304 | 572 | 570 | 0.873 | 0.788 |
| 5-TAMRA | -0.561 | -0.505 | -0.654 | -0.619 | 591 | 592 | 0.862 | 0.799 |
| Alexa 594 | 0.181 | 0.086 | -0.889 | -0.848 | 635 | 626 | 0.907 | 0.853 |
| NADH | -0.305 | -0.151 | 0.440 | 0.474 | 504 | 498 | 0.536 | 0.498 |
| FAD | -0.718 | -0.642 | -0.172 | -0.048 | 561 | 556 | 0.738 | 0.644 |

**Fluorescent dye solutions measured with the sine/cosine filter method compared to 32 channel spectral detection.** Table comparing the S, G, emission wavelength center of mass, and modulation obtained with both methods. Please note that the G and S coordinates are not directly comparable due to the slightly different detection ranges of the two methods (sine/cosine: 400-700 nm, 32 channel detector: 410-695 nm). The wavelength and modulations account for those ranges and are comparable. Two-photon excitation (800 nm) was used to avoid instrument-specific effects of notch filters and dichroic mirrors in the light paths.

**Supplementary Fig. S3.**

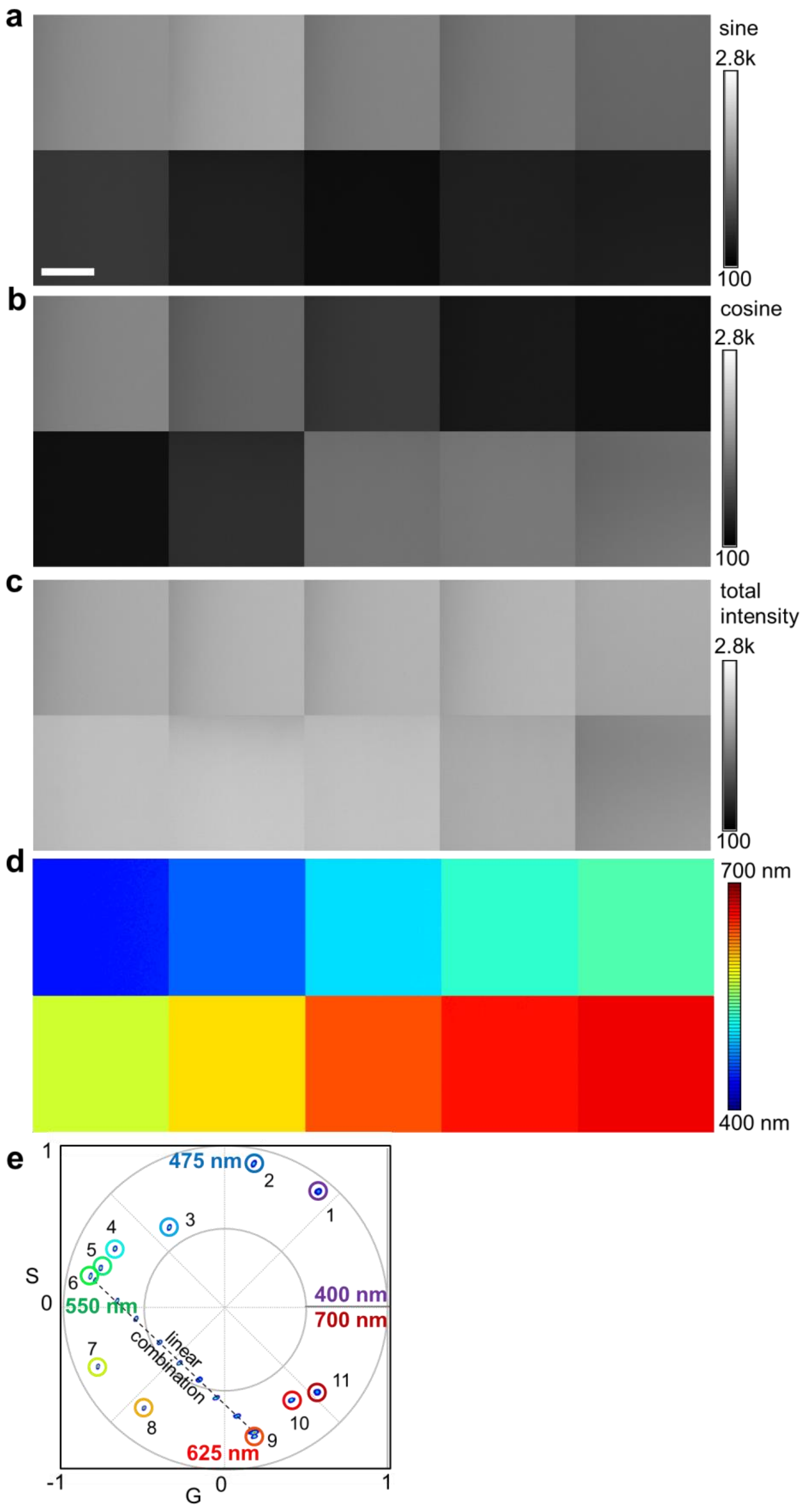

**Raw images and phasor plot histograms of eleven dye solutions and demonstration of the rule of linear addition.** (**a-c**) Collages of the raw sine, cosine, and total intensity images of the eleven dye solutions with fluorescence emissions from blue to deep red (left to right, top to bottom). (**d**) Dye solution collage color coded according to the selection on (**e**) the phasor plot; (1) 9-10 Diphenyl-anthracene, (2) Coumarin 1, (3) ANS, (4) Coumarin 6, (5) Fluorescein, (6) Rhodamine 110, (7) Rhodamine 6G, (8) 5-TAMRA, (9) Atto 590, (10) Nile Red, and (11) MitoTracker Deep Red. In addition, the law of linear addition was demonstrated by mixing Rhodamine 110 and Atto 590 at different proportions (dashed line). Image data shown represents an average of 50 frames of 256 x 256 pixels. No emission filters other than sine/cosine filters and a 400-700 nm bandpass were used. Scale bar 10 µm.

**Supplementary Fig. S4.**

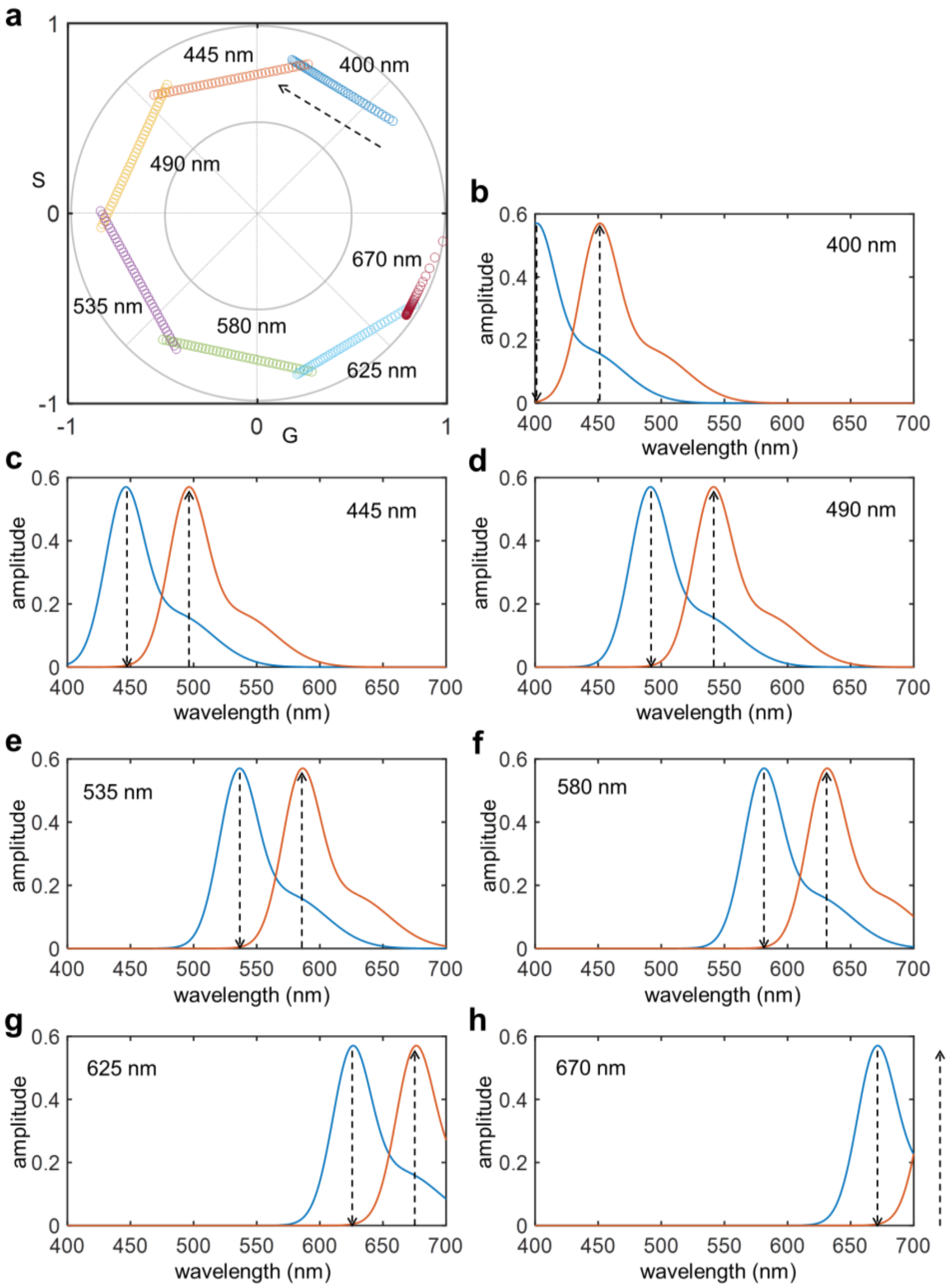


**Phasor plot rule of linear addition demonstrated by simulations.** Emission spectra of two fluorophores were simulated with their peak emissions separated by 50 nm. Their relative amplitudes were ramped from 100% to 0% for the blue shifted peak and 0% to 100% for the red shifted peak in 2.5% steps. The simulation was repeated seven times with the first peak starting at 400 nm in 45 nm increments. (**a**) Phasor plot of the linear combinations of the seven combinations. The arrow indicates the direction of the amplitude ramp, the wavelengths indicate the peak emissions of the blue shifted peak for each simulation. (**b-h**) Simulated emission spectra used for the phasor plot transformations at the 50% amplitude points with the amplitude ramps indicated by arrows. It can be seen that the linearity (rule of linear addition) is always maintained. The peak ratios as determined by phasor analysis become unevenly spaced as the area under the emission curve is significantly clipped for the peaks at the blue and red edges of the simulated detection range of 400-700 nm (b, g, h). This could be accounted for in experiments by intensity calibrations with the pure species of known (relative) concentrations.

**Supplementary Fig. S5.**

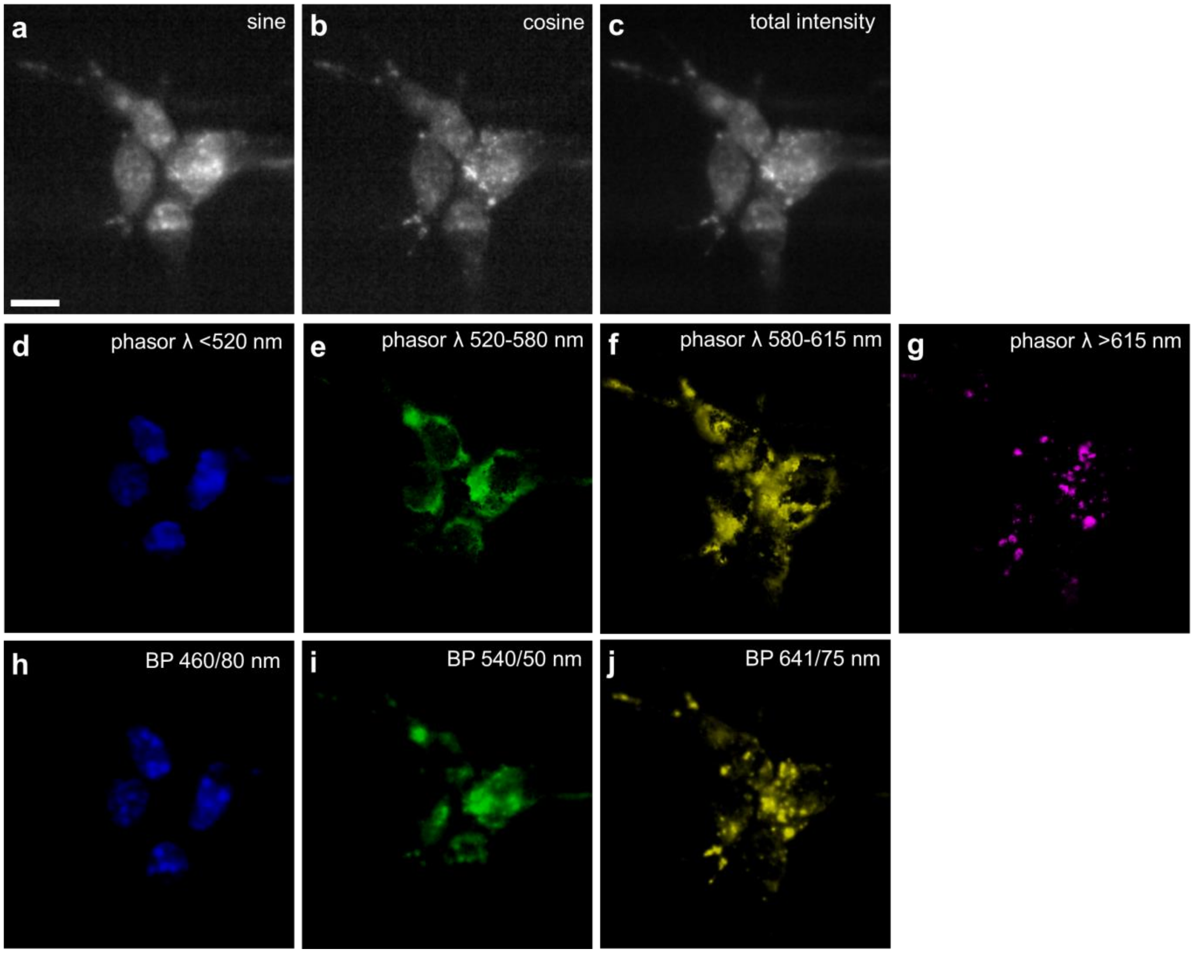


**Raw sine, cosine, total intensity, unmixed phasor and bandpass filter images of live cells stained with four dyes.** NIH 3T3 cells were stained with a mixture of NucBlue, BODIPY FL, MitoTracker Orange, and LysoTracker Red and subjected to hyperspectral light sheet imaging with sine/cosine filters as well as three bandpass filters (DAPI/FITC/TexasRed). (**a-c**) Raw camera images acquired with sine, cosine, and without filter. (**d-g**) Voxel center emission wavelength selection according to the indicated phasor λ ranges. (**h-j**) Bandpass filter images in the indicated spectral windows are not able to accurately separate the colors and separation of mitochondria and lysosomes is completely absent. Sample was excited with 870-nm light *via* two-photon absorption. Scale bar, 10 µm.

**Supplementary Fig. S6.**

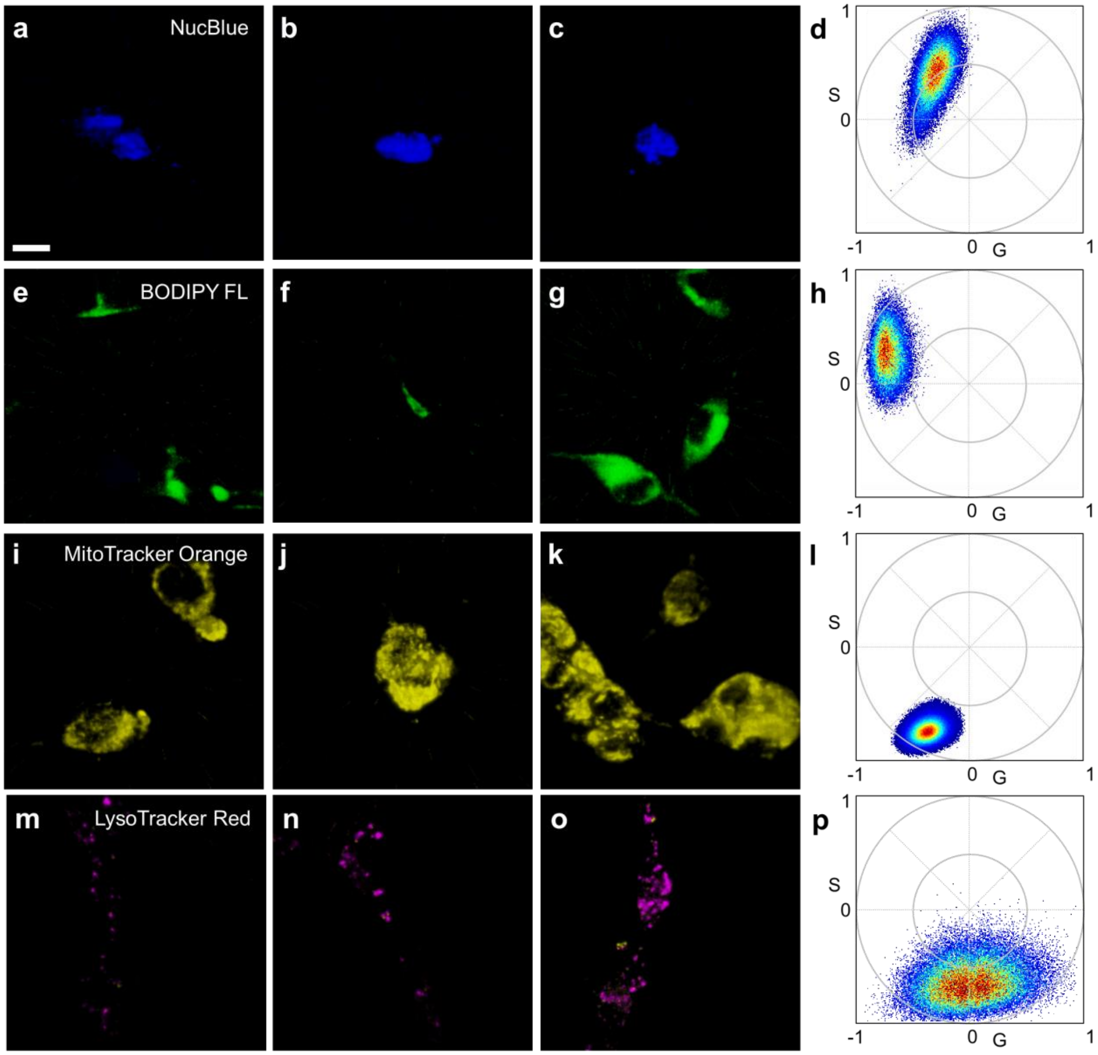


q

| Cell Stain | G | S | λ (nm) | Modulation |
|---|---|---|---|---|
| NucBlue | -0.3236 | 0.3836 | 511.5229 | 0.5558 |
| Bodipy FL | -0.726 | 0.2861 | 533.1365 | 0.8097 |
| MitoTracker Orange | -0.3889 | -0.7331 | 601.9121 | 0.8359 |
| LysoTracker Red | 0.0303 | -0.6142 | 623.9926 | 0.7288 |

**Single stained NIH 3T3 cells and corresponding phasor plot distributions.** NIH 3T3 cells were stained with either (**a-d**) NucBlue, (**e-h**) BODIPY FL, (**i-l**) MitoTracker Orange, or (**m-p**) LysoTracker Red and subjected to hyperspectral light sheet imaging with sine/cosine filters. Shown are the color coded fluorescence volumes and the corresponding phasor plots. (**q**) Average *G*, *S*, center emission wavelengths, and modulations. Samples were excited with 870-nm light *via* two-photon absorption. Scale bar, 10 μm.

**Supplementary Fig. S7.**

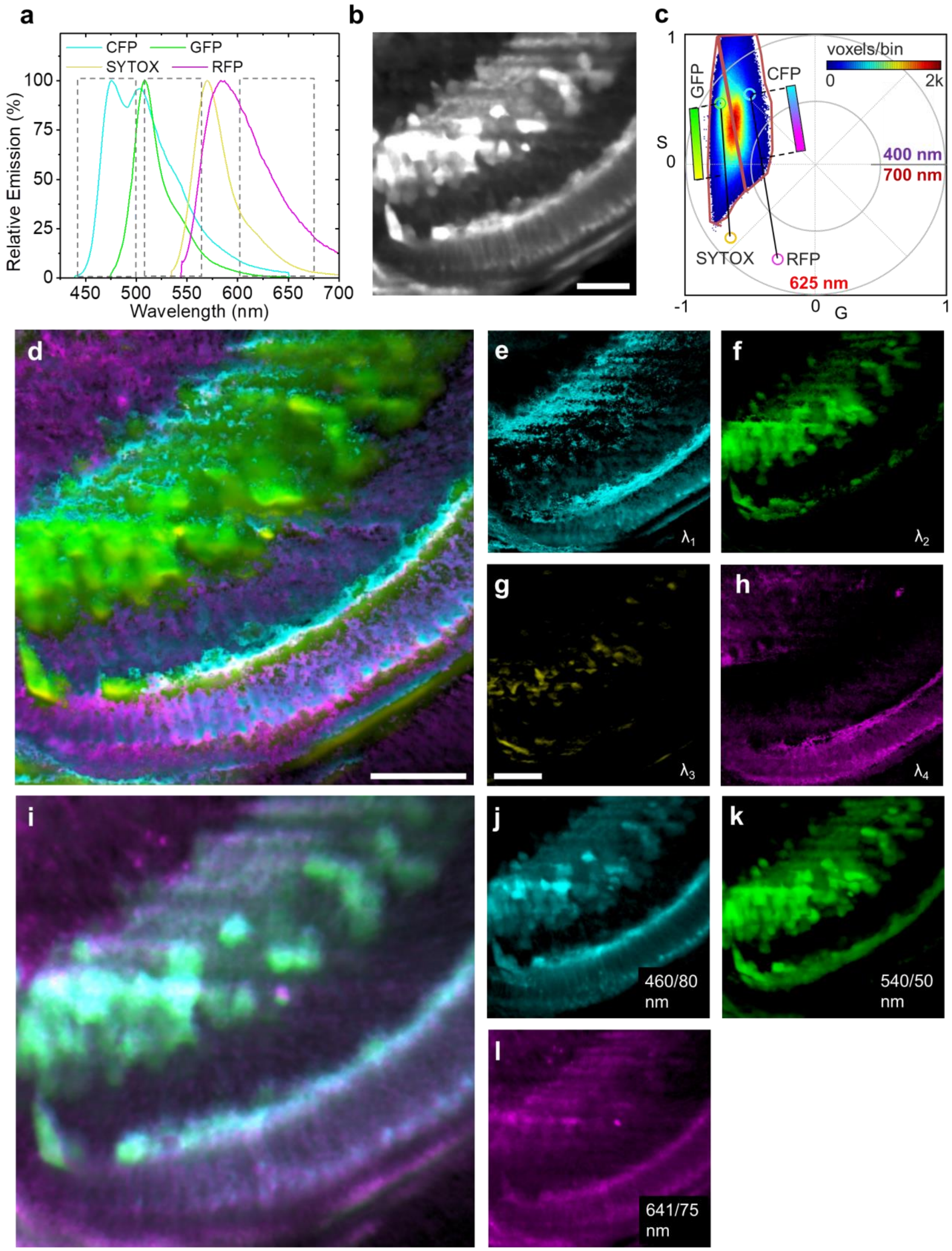


**SoFa transgenic 72 hpf zebrafish imaged with conventional bandpass filters and the sine/cosine hyperspectral method.** SoFa transgenic zebrafish were fixed and stained with

SYTOX ORANGE in addition to the labeled constructs Crx:gapCFP, Ptf1a:cytGFP, and Atoh7:gapRFP. (**a**) Emission spectra and bandpass filter windows (spectra raw data, Invitrogen). (**b**) Fluorescence intensity light sheet image collected in a 400-700 nm window. (**c**) Phasor plot obtained by the sine/cosine filter method of 21 *z* slices (1 µm steps). Circles indicate the theoretical phasor plot positions of pure CFP, GFP, SYTOX, and RFP. The phasor histogram was divided into two halves, the voxels in each half were color coded according to their position along the connecting lines of the two fluorophore pairs (CFP/RFP, GFP/SYTOX) with the indicated color maps (cyan/magenta, green/yellow). (**d**) Merged and (**e-h**) individual phasor color mapped images. (**i**) Merged and (**j-l**) individual images of the same volume imaged with bandpass filters as indicated in (a). Clearly, with bandpass filters, separation of CFP/GFP signals was not possible. While the instrument was equipped with three bandpass filters (six position filter wheel: total intensity, sine, cosine, DAPI, FITC, TxRed), the addition of a forth bandpass filter would not have helped to separate RFP/SYTOX Orange fluorescence as their overlap is even stronger than the spectral overlap between CFP and GFP. Scale bars, 25 µm.

**Supplementary Fig. S8.**

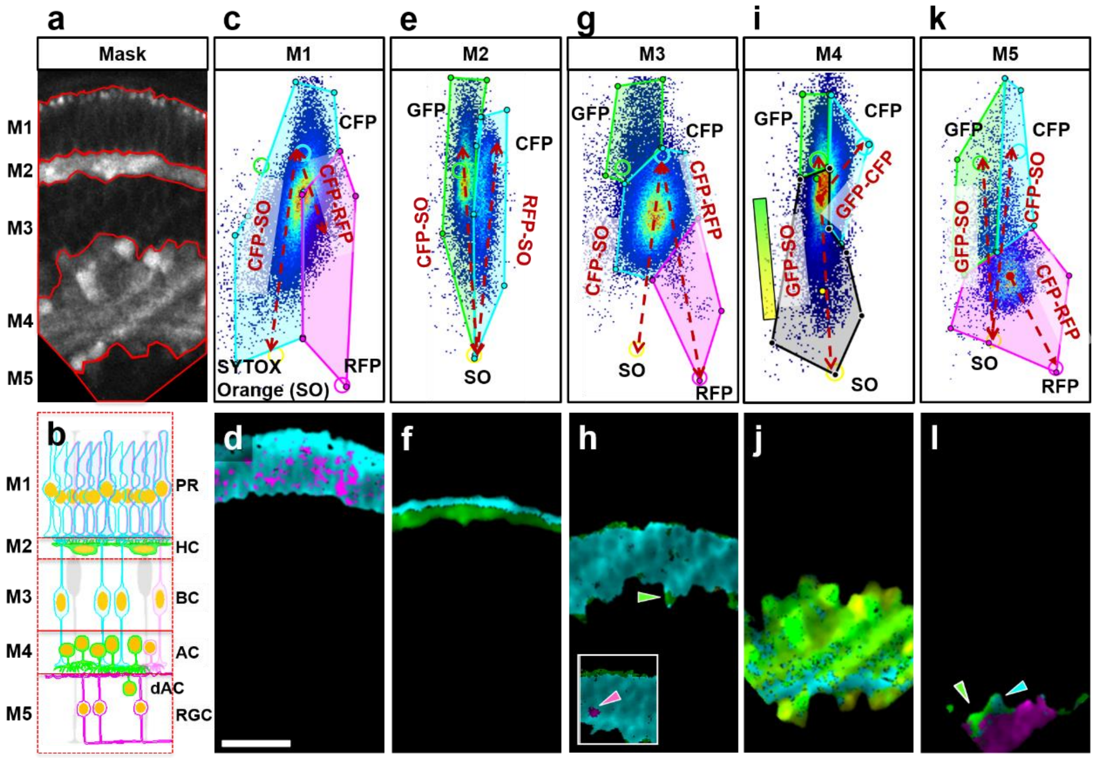


**Hyperspectral cellular fingerprinting reveals zebrafish retinal composition.** (**a**) Fluorescence intensity image showing the region of the retina used for segmentation phasor analysis; red lines indicate masks (M1-M5) for the cell layers of the SoFa fish retina. (**b**) Scheme of the different cell types and layer organization of the zebrafish retina at this embryonic stage. (**c**) M1 phasor plot identifying the linear combination between CFP and SYTOX Orange (SO) (labeled CFP-SO) and a group of voxels with a fraction of CFP/SO plus RFP (labeled CFP-RFP). (**d**) Pseudocolored image representing the voxels selected inside the polygons. The pattern corresponded well with the expected high expression of CFP and low expression of RFP in the PR. (**e**) M2 phasor plot identifying two linear combinations, CFP-SO and GFP-SO. Polygon voxel selections highlight (**f**) CFP expressed in the upper boundary of the HC layer in addition to strong GFP expression from HC. (**g**) M3 phasor plot showing a big voxel cluster at the center between all components, associated with a linear combination of CFP and SO (labeled CFP-SO) marked with the cyan polygon. (**h**) The few voxels selected by the green polygon identified an amacrine cell (green arrowhead) of the adjacent layer (HC), showing the high spectral sensitivity of the phasor plot in each voxel. Additional weak expression of GFP and RFP was detected in this layer as well (green and magenta polygons), visible in some areas (inset, magenta arrowhead). (**i**) M4 phasor plot identifying the linear combination of GFP and SO highlighted with a black polygon and colored with a gradient color scheme (green-yellow) labeled as GFP-SO. This layer included a small group of voxels towards the CFP location, identified with the cyan polygon. (**j**) The pseudocolored image showed that the GFP-SO gradient corresponded well to the expected expression of GFP in amacrine cells, together with the nuclear SO stain. The cyan region highlighted a group of voxels at the bottom of the pseudocolored image, associated with axons of BP cells expressing CFP. (**k**) M5 phasor plot identifying a main cluster of voxels associated with a linear combination of RFP/SO highlighted by the magenta polygon. Two more groups were observed in this layer with either GFP or CFP and different fractions of SO (green and cyan polygons). (**l**) The pseudocolor image revealed that the magenta polygon selection corresponded to RGC, while the green polygon selection was consistent with displaced amacrine cells (dAC) (green arrowhead), and the cyan polygon selection with BP neurites (cyan arrowhead). Scale bar, 25 µm.

**Supplementary Fig. S9.**

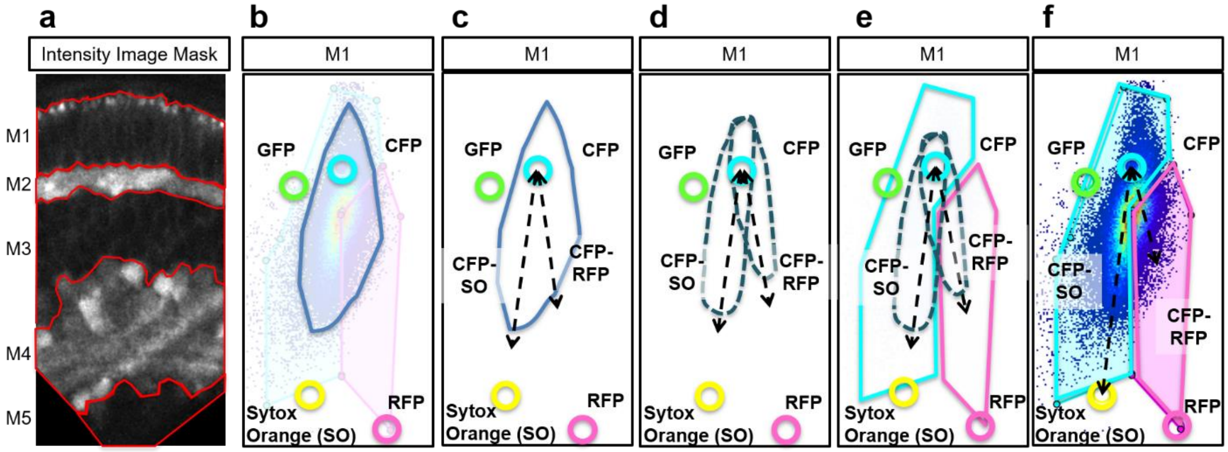


**Principle of phasor plot analysis based on linear combinations.** (**a**) In a first step, taking advantage that retinal layers are evident at this stage, the total intensity images were used to segment the image into regions of interest (M1-M5), using anatomical criteria (**see Fig. S6b**). In the following, the M1 photoreceptor (PR) layer was selected and used as an example. (**b**) The phasor plot distribution of pixels in the selected M1 region is shown as 2D histogram, higher voxel densities are indicated by a color scheme ranging from blue to green to yellow and to red. The cyan, green, yellow and magenta circles indicate the theoretical positions of the pure fluorophores (CFP, GFP, SO, RFP) on the phasor plot. A blue line outlines the perimeter of the voxel distribution. (**c**) The shape of this particular distribution is consistent with the existence of two linear combinations in the M1 region: CFP-SO and CFP-RFP, indicated by dashed arrows. This is further illustrated in (**d**), where dashed outlines represent the two clouds expected for an image with such underlying linear combinations. (**e**) To highlight the two linear combinations, we selected the areas indicated by cyan and magenta polygons. Panel (**f**) shows the actual phasor plot data superimposed with the areas selected. The colors (here cyan and magenta) of the selected voxels are then remapped to the intensity image shown in **Fig. S6d**.

**Supplementary Fig. S10.**

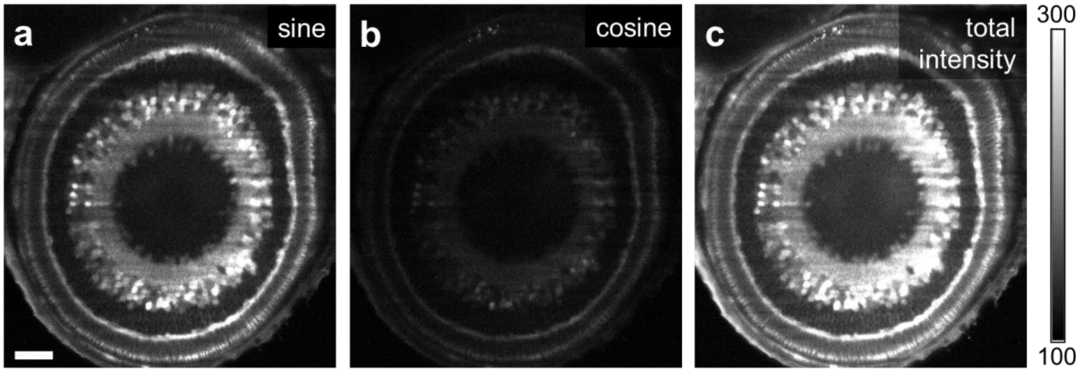


**Raw sine, cosine, and total intensity images of a SoFa zebrafish retina.** Single *z* section of unprocessed raw camera images as obtained with the sideSPIM through the (**a**) sine, (**b**) cosine, and (**c**) without filter. Scale bar, 50 μm.

**Supplementary Fig. S11.**

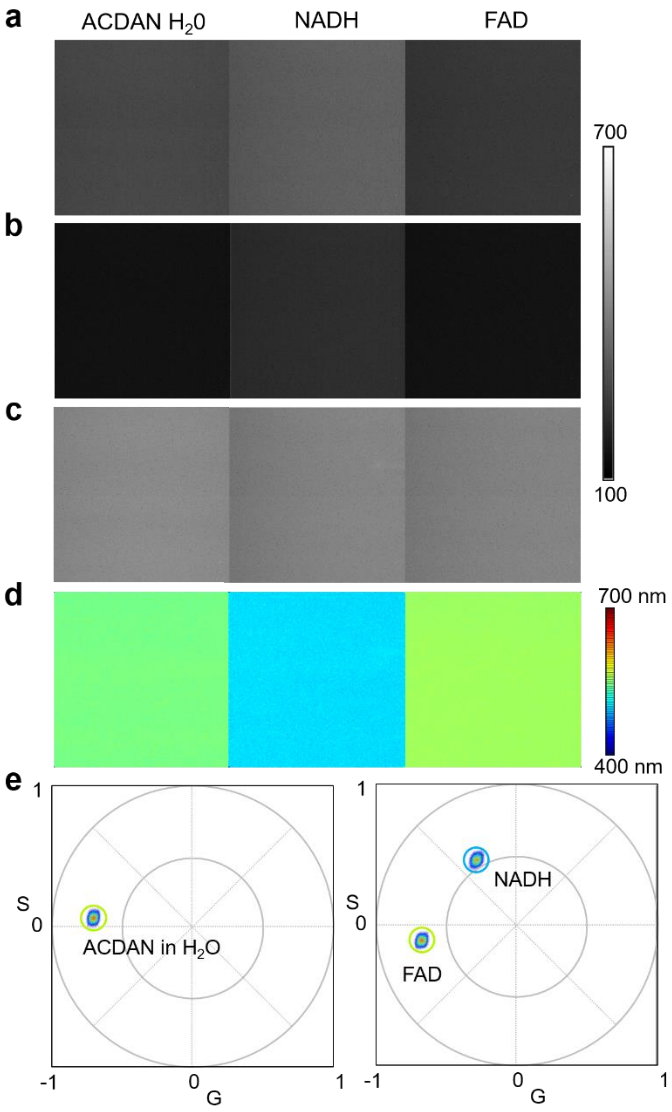


**Spectral phasor positions of ACDAN, NADH and FAD in aqueous solutions.** A solution of ACDAN (30 μM) was prepared in nanopure water ($H_2O$) and subjected to sine/cosine imaging. Solutions (2 mM) of NADH and FAD were prepared in PBS buffer. Fluorescence was excited at 740 nm (NADH), 780 nm (FAD), and 800 nm (ACDAN) by two-photon excitation. (**a-c**) Raw sine, cosine, and total intensity images. (**d**) Images color coded according to the center wavelengths of emission. (**e**) Phasor plot representations of the data as 2D histograms. Scale bar, 10 μm.
