## Supplementary figures and images for "Ultrafast phasor-based hyperspectral snapshot microscopy for biomedical imaging"

### Movie M1

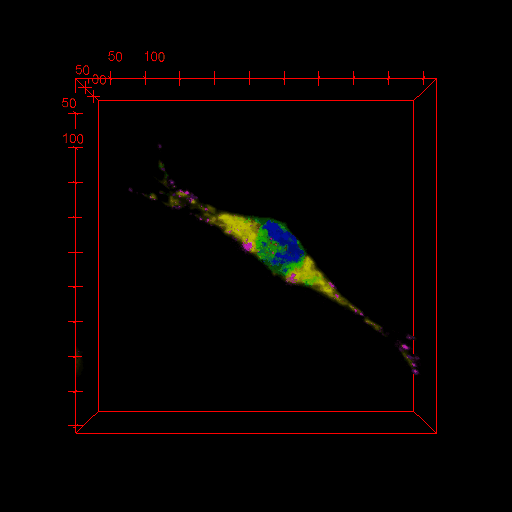

### Movie M2

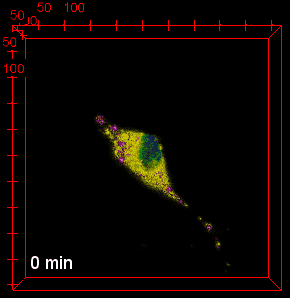

### Movie M3

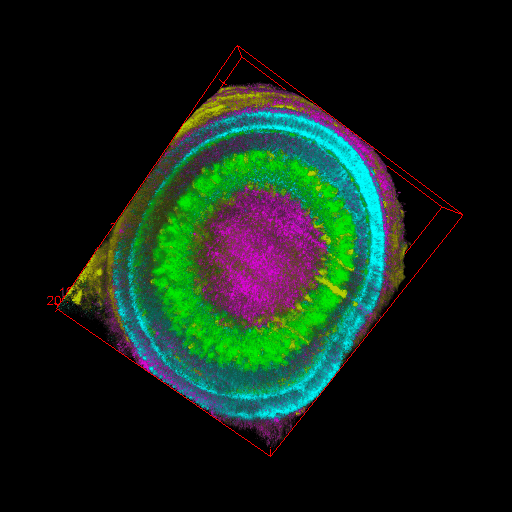
